# BiomeGPT: A taxonomy-aware foundation model for the human microbiome

**DOI:** 10.64898/2026.01.05.697599

**Authors:** Nicholas A. Medearis, Nicholas S. Yousefi, Ali R. Zomorrodi

## Abstract

The human microbiome encodes rich information about host health, yet current analysis pipelines remain narrowly optimized for individual tasks. This limits our ability to gain a thorough view of how the microbiome impacts health and disease. Here, we introduce BiomeGPT, a taxonomy-aware microbiome foundation model pretrained on 13,500 human metagenomes spanning 34 phenotypes, including healthy and 33 diseases, to learn species-level representations of microbiome composition. BiomeGPT explicitly encodes within-sample species abundance patterns and incorporates the full microbial taxonomic hierarchy into species representations, encompassing all eight taxonomic ranks. This enables the model to capture fine-grained microbial community composition, taxonomic relationships among species, and context-dependent cross-species dependencies embedded within community profiles. When fine-tuned for predicting host health status, BiomeGPT accurately distinguishes healthy from diseased microbiomes and resolves individual disease types across a broad clinical spectrum. Furthermore, differential attention analysis showed that phenotype-associated shifts in model attention often aligned with differential abundance, but the strength and direction of this relationship varied markedly across diseases, indicating that the model captures signals beyond univariate abundance changes alone. By providing a unified, scalable framework for microbiome representation learning and prediction, BiomeGPT enables new avenues for investigating microbiome–host phenotype relationships across diverse clinical contexts.

## Introduction

The human microbiome is a complex and dynamic ecosystem that plays an important role in health and disease through extensive interactions with host physiology. Among microbial communities across different body sites, the gut microbiome, in particular, contributes to a broad range of host physiological processes, including digestion of complex nutrients, modulation of immune and metabolic functions, and maintenance of intestinal homeostasis. Disruptions in the gut microbiome’s composition and function—commonly referred to as gut dysbiosis—have been linked to a broad range of conditions, including gastrointestinal conditions like inflammatory bowel disease (IBD) and colorectal cancer, systemic conditions such as obesity, type 2 diabetes, cardiovascular disease, and neurodegenerative disorders ^1, 2^. Due to its far-reaching effects on human physiology and disease, the microbiome has emerged as a prime area of investigation for uncovering disease mechanisms and developing new therapies. Because the microbes within the microbiome function as a tightly connected network rather than isolated players, a holistic, systems-level perspective is essential to unravel how shifts in community structure translate into health or disease.

Over the past decade, the computational toolkit for microbiome data analysis to characterize microbiome alterations and their association with host health has significantly expanded, with a wide array of tools developed for differential abundance analysis ^3, 4^, multivariate analysis ^5^, biomarker discovery ^6^, network inference ^7^, and microbiome-based prediction of host phenotypes ^8–10^, among others. Despite this methodological breadth, existing computational tools are rather scattered, each specializing in a specific analytical task. Likewise, the datasets used for each tool are typically task-specific and limited in scale and scope. As a result, microbiome studies often rely on ad hoc combinations of independent pipelines, leading to fragmented analytical workflows, with limited ability to transfer information learned from one cohort, phenotype, or task to another. This compartmentalized approach can also hinder the identification of shared and context-dependent relationships across heterogeneous microbiome datasets, such as how variations in microbial community structure influences clinical outcomes and disease risk across diverse phenotypes and cohorts. Capturing such cross-task relationships is essential for advancing a systems-level understanding of the complex interactions within microbiomes and between microbiomes and human hosts.

Generative AI and transfer learning have transformed fields such as natural language processing, by enabling the development of general-purpose foundation models that can be adapted to diverse tasks through a two-stage process: pretraining on massive unlabeled datasets, followed by fine-tuning for specific tasks. This paradigm offers substantial advantages in transferability, scalability, and performance. Inspired by these advances, a key question emerges for microbiome research: could a similar framework support a unified representation-learning approach for microbiome-based modeling and prediction across diverse phenotypes, rather than relying on separate, task-specific models?

Foundation models have already begun to reshape life science research. A series of foundation models have been developed recently across multiple domains of biology, each pretrained on massive biological data and then fine-tuned to specific tasks. In structural biology, AlphaFold series of models ^11–13^ have set a new standard by accurately predicting protein 3D structures from sequence alone. RNA-focused models such as RhoFold+ ^14^ and RNA-FM ^15^ have extended this paradigm to RNA, showing that large-scale pretraining on millions of RNA sequences enables accurate prediction of RNA 3D structure, secondary structure, and interhelical geometry directly from sequence. In protein modeling, large-scale generative language models such as ProGen ^16^ have proven capable of learning the sequence–function relationships. In single-cell transcriptomics, foundation models such as scBERT ^17^, Geneformer ^18^, and scGPT ^19^ have been trained on tens of millions of scRNA-seq profiles, capturing the high-dimensional structure of cellular expression data. These models illustrate the power of transfer learning, shared representations, and generative architectures to unify diverse tasks across biology.

These advances have largely focused on molecular and cellular levels of biological organization. The microbiome presents a distinct modeling challenge at the community level, where host-associated phenotypes emerge in the context of complex interactions among microbial taxa and their collective community structure, highlighting the need for foundation models specifically designed for microbiome data. A number of microbiome-specific foundation models ^20–22^ have already demonstrated the potential of large-scale pretraining in this domain. However, these models largely rely on ASV-or genus-level taxonomic representations from amplicon sequencing data or rank-based tokenization schemes and represent microbial taxa as independent vocabulary elements without explicitly accounting for their evolutionary relationships.

In this study, we sought to develop a microbiome foundation model that learns fine-grained representations of microbial community composition while explicitly incorporating taxonomic hierarchy. We introduce BiomeGPT, a species-level foundation model pretrained on large-scale metagenomic profiles spanning a broad range of clinical conditions. We evaluated whether the learned representations could be transferred across diverse host health and disease prediction tasks and examined phenotype-associated microbial patterns through differential attention analysis. Together, these analyses establish a framework for learning reusable, taxonomy-aware representations of human microbiomes across diverse health and disease contexts.

## Results

To evaluate the potential of a foundation model for microbiome representation learning, we developed BiomeGPT, a transformer-based architecture pretrained on taxonomy-aware species-level abundance profiles from 13,500 human metagenomes spanning multiple body sites and diverse clinical conditions. BiomeGPT integrates quantitative abundance information with taxonomic hierarchy and uses masked abundance prediction to learn representations of microbial community composition. We first evaluated the biological structure captured by the pretrained representations and then fine-tuned the model across a range of host health prediction tasks, including classification of healthy vs. diseased samples and differentiation across 26 disease types. We further used differential attention analysis to characterize microbial species associated with host health and disease phenotypes and to evaluate the relationship between differential attention and differential abundance.

### Pretraining and taxonomy-aware representation learning on human metagenomes

#### Dataset overview

To train BiomeGPT, we assembled a large dataset of human metagenomic samples curated from three publicly available resources: the curatedMetagenomicData (cMD) repository ^23^, the Gut Microbiome Health Index (GMHI) ^24^, and the Gut Microbiome Wellness Index 2 (GMWI2) ^25^. All samples in these resources were taxonomically profiled at species-level resolution using MetaPhlAn ^26, 27^, ensuring a unified and consistent species-level feature space across all samples and studies within the entire training dataset. For longitudinal studies, only the sample from the initial time point for each subject was retained. After filtering, the training dataset included 13,524 metagenomes, predominantly from gut/stool samples but also including samples from other body sites, spanning healthy and disease-associated microbiomes. Following species-level preprocessing (see **Methods**), 13,500 samples were retained for pretraining. The training dataset was drawn from 85 studies, encompassed 34 phenotypes (healthy and 33 diseases), and included 1,028 unique microbial species.

We further created a gut-specific dataset by filtering out non-gut/stool samples representing five phenotypes (Mucositis, Periodontitis, Peri-implantitis, Psoriasis, Arthritis), along with associated species from the training dataset. The resulting gut-specific training dataset included 13,333 samples, 895 species, and 30 phenotypes (including healthy and 29 diseases). The full pretraining datasets are available in **Supplementary File 1**.

We additionally created an external validation dataset consisting of 927 samples from 12 studies. To enable rigorous evaluation of model generalization, all samples present in the external validation set came exclusively from studies not represented within the training set. The validation dataset contained only gut/stool samples and spanned healthy and five disease phenotypes, including Ankylosing Spondylitis, Colorectal Cancer, Crohn’s disease, Parkinson’s disease, and Rheumatoid Arthritis. A breakdown of sample counts per disease for the gut-specific training and external validation datasets is provided in **Fig. 1**.

**Fig. 1.**
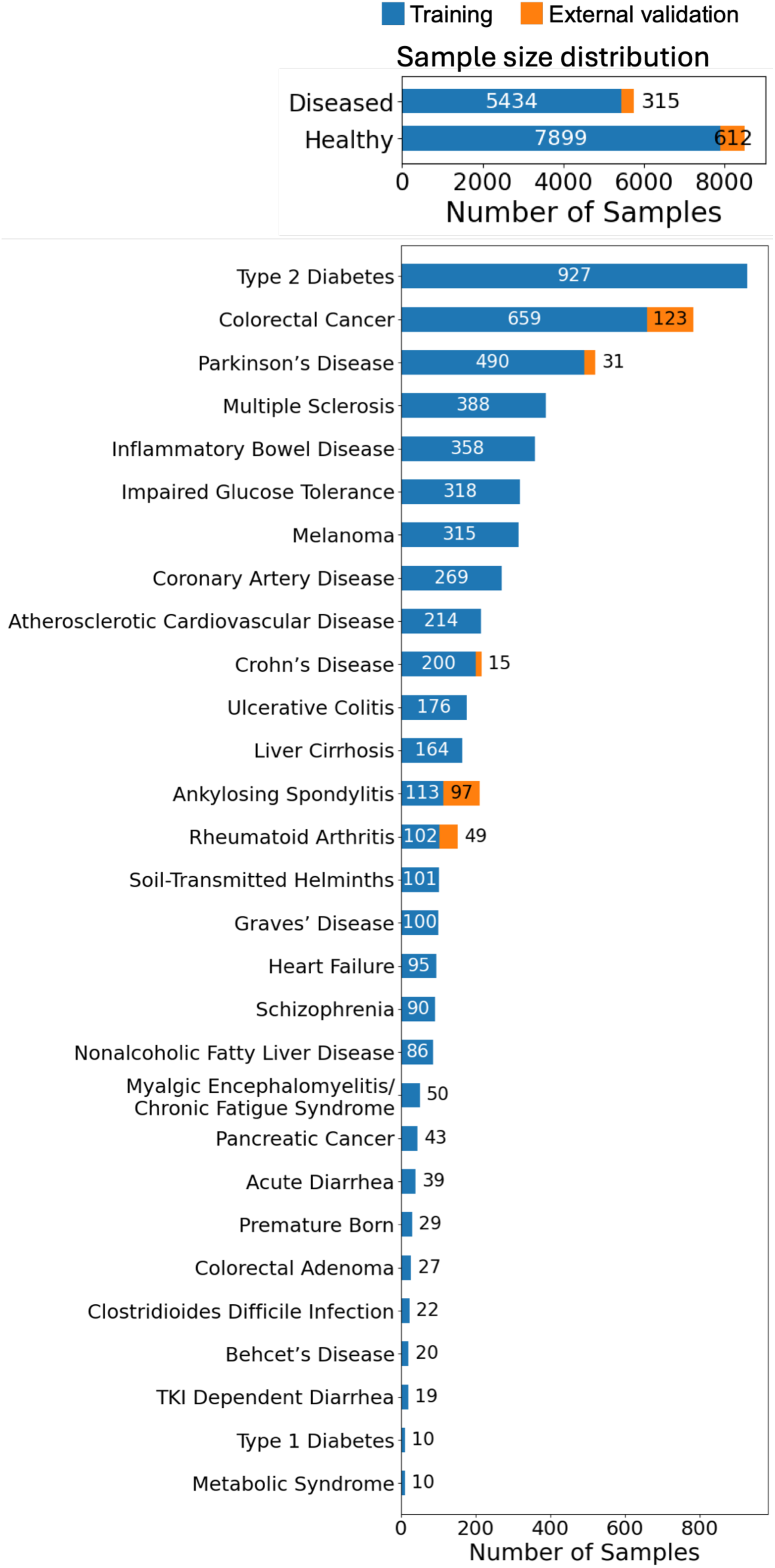
Overview of the gut metagenomic datasets used for pretraining and external validation of BiomeGPT. Number of samples per phenotype, separated by training and external validation cohorts, is shown for all phenotypes with 10 or more samples.

#### Abundance binning

To prepare species-level taxonomic profiles for pretraining, we converted continuous relative abundances (from MetaPhlAn ^26, 27^) into discrete abundance bins. This transformation—adapted from discretization strategies used in models such as scBERT ^17^ and scGPT^19^—captures the relative magnitude of species abundance within each sample. Specifically, species were sorted by abundance and assigned to one of the *B* bins, with undetected species (i.e., those with zero relative abundance) assigned to bin 0 (see **Methods**). This quantile-based approach converts continuous abundances into discrete within-sample abundance states, encoding coarse abundance information explicitly for each species rather than using its position in an abundance-ranked sequence. It may also reduce sensitivity to absolute abundance scale and some study-specific technical variation arising from differences in sequencing depth, sequencing platform, or library preparation. The resulting binned abundance matrix served as the model input for masked modeling during pretraining.

#### Taxonomy-aware embedding strategy and microbiome sample tokenization

To model gut microbiomes using a transformer-based architecture, we represented each sample as a sequence of microbial species together with their binned abundance values, yielding a set of (species, abundance) pairs that served as input tokens for the model. Instead of representing microbial species as independent vocabulary elements, each species was represented using a taxonomy-aware embedding that incorporates its position within the microbial taxonomic hierarchy. Specifically, learnable embeddings were assigned across the eight taxonomic ranks, from Domain to Species, and combined to generate the species-level representation (**Fig. 2**). This allows related species to share information through their common taxonomic ancestry while retaining species-specific identity. Each discrete abundance bin was independently mapped to an embedding through a lightweight multilayer perceptron (MLP). The species and abundance embeddings were normalized separately using layer normalization to place them in comparable representational spaces and then combined additively to generate the final input token for each (species, abundance) pair (**Fig. 2**).

**Fig. 2.**
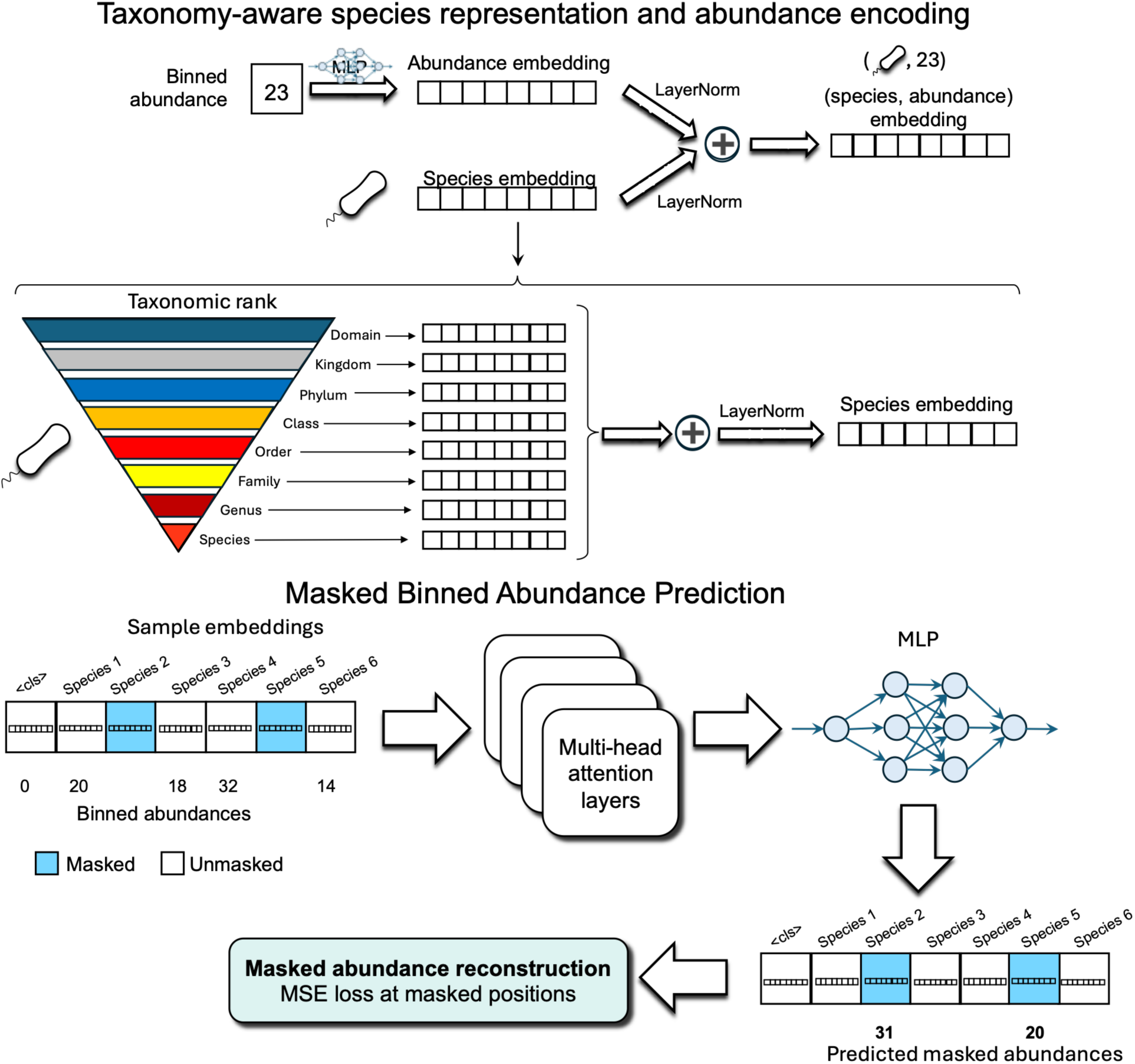
Overview of the taxonomy-aware embedding strategy and pretraining architecture of BiomeGPT. Each microbiome sample is represented as a sequence of species and binned abundance pairs. Species embeddings incorporate taxonomic hierarchy across the eight taxonomic ranks and are combined with abundance embeddings to generate input token representations. During pretraining, a subset of species with nonzero abundance is randomly masked, and masked attention allows information flow among unmasked species and from masked species to the unmasked community context and themselves (also see **Supplementary Fig. 1**). The resulting representations are processed through multi-head attention layers and an MLP to predict the binned abundances of masked species, with mean squared error computed at masked positions.

To enable downstream classification, we introduced a special `<cls>` token at the beginning of each sample sequence to represent sample-level context (e.g., healthy or diseased). This token had a dedicated learnable embedding and was always assigned an abundance bin value of 0. The `<cls>` token serves as a placeholder for aggregating information across the entire microbiome sample through attention.

#### Transformer architecture and pretraining strategy

BiomeGPT employs a transformer-based architecture to learn latent representations of microbiomes through masked abundance prediction. During pretraining, we used a masked attention strategy tailored to the unordered nature of microbial species. In each sample, a random subset of species was masked, and the model was trained to reconstruct their binned abundance values from the unmasked community context. Attention was restricted to occur between unmasked species and from masked species to the unmasked species and themselves (see **Methods** and **Supplementary Fig. 1),** enabling the model to learn species co-occurrence and abundance relationships without imposing an arbitrary positional structure. Species with zero abundance were excluded from attention and loss computation. Pretraining comprised two complementary objectives adapted from the gene-prompt and cell-prompt strategies in scGPT ^19^. Both objectives reconstruct masked species abundances from microbiome context, with the species prompt emphasizing species-to-species relationships and the sample prompt encouraging the `<cls>` token to capture sample-level community structure. The overall pretraining loss combines the reconstruction losses from both objectives (see **Methods**). BiomeGPT was pretrained in two phases. The model was first trained on 13,500 human metagenomic samples spanning gut and non-gut body sites, followed by further pretraining on the gut-specific dataset of 13,333 samples to specialize the learned representations for the intestinal microbiome.

#### Unsupervised evaluation of sample-level representations learned by BiomeGPT

To assess whether BiomeGPT learns biologically meaningful structure prior to supervised fine-tuning for health status prediction, we evaluated the pretrained model’s sample-level representations using the `<cls>` token embeddings derived from both gut and non-gut metagenomic samples (i.e., before gut adaptation). Low-dimensional visualization using UMAP revealed a separation between gut and non-gut samples in the learned embedding space (**Fig. 3A**), indicating that body-site information is captured at the community level without explicit supervision. This indicates that BiomeGPT encodes robust, body-site– specific structure in its sample-level representations, supporting the model’s ability to learn generalizable microbiome features during pretraining. Of note, a small subset of gut samples formed a distinct region of the embedding space; these samples were primarily drawn from disease-only cohorts and exhibited distinct taxonomic composition, consistent with disease- and cohort-associated structure rather than non-gut contamination.

**Fig. 3.**
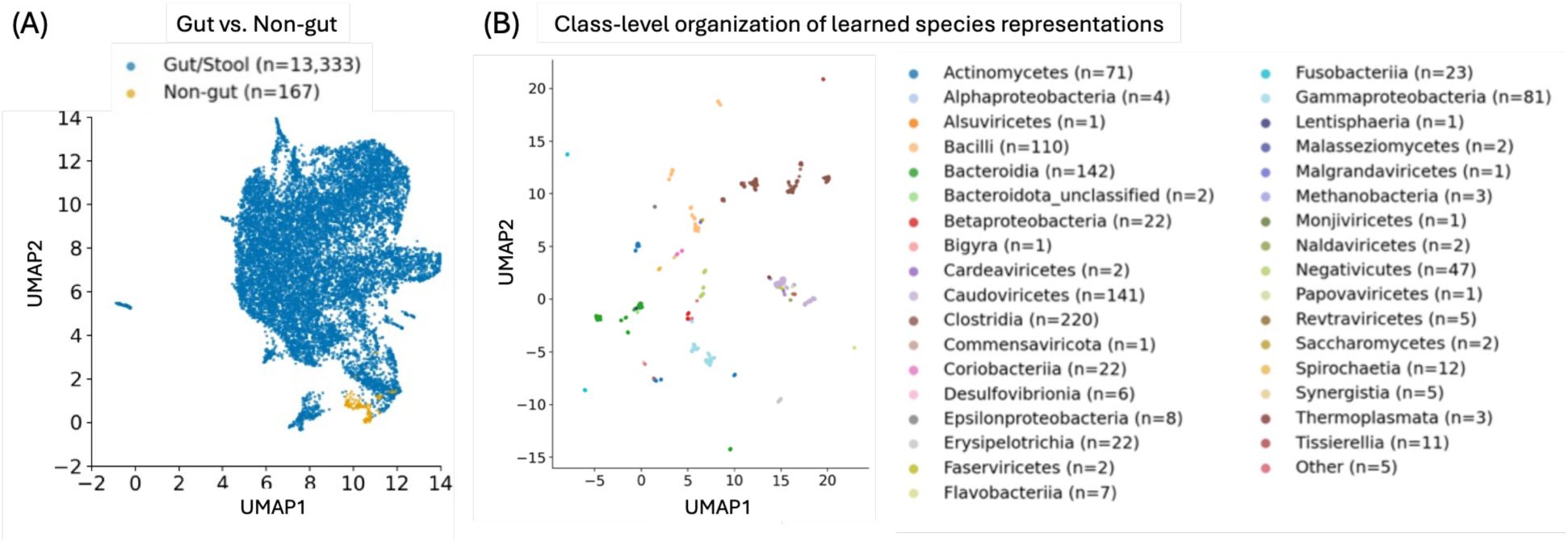
Unsupervised evaluation of sample- and species-level representations learned by BiomeGPT during pretraining. (A) UMAP visualization of sample-level `<cls>` representations of gut/stool and non-gut microbiome samples. (B) UMAP visualization of learned species representations colored by taxonomic class (n = 621), showing preservation of taxonomic structure in the learned representation space.

Visualization of the learned species embeddings further showed that the taxonomy-aware representation preserved hierarchical taxonomic structure. At the class level, species belonging to the same taxonomic class tended to occupy coherent regions of the embedding space (**Fig. 3B**), with similar organization also observed at the order and phylum levels (**Supplementary Fig. 2**). These patterns indicate that the learned species representations retain taxonomic relationships across multiple levels of the microbial hierarchy.

### Fine-tuning and evaluation of BiomeGPT for health status prediction across diverse disease phenotypes

Following pretraining, we evaluated the utility of BiomeGPT for supervised clinical inference by fine-tuning the model on a range of classification tasks designed to infer host health status from gut microbiome profiles. Specifically, we explored classification tasks including Healthy vs. Diseased and Individual Disease vs. Healthy across diverse diseases.

#### Dataset preprocessing

Before fine-tuning, we further preprocessed the gut-specific dataset to reduce potential contamination and retain species robustly represented across phenotypes. This included removal of predefined non-gut or environmental taxa and phenotype-specific prevalence and abundance filtering (see **Methods**). The resulting dataset contained 13,333 samples and 621 distinct species (**Supplementary File 1**).

#### Model architecture

For all tasks, we appended a downstream classification head to the pretrained model and applied the same model architecture. The learned sample-level `<cls>` token from the transformer was provided as input to this classifier (see **Fig. 4A** and **Methods**).

**Fig. 4.**
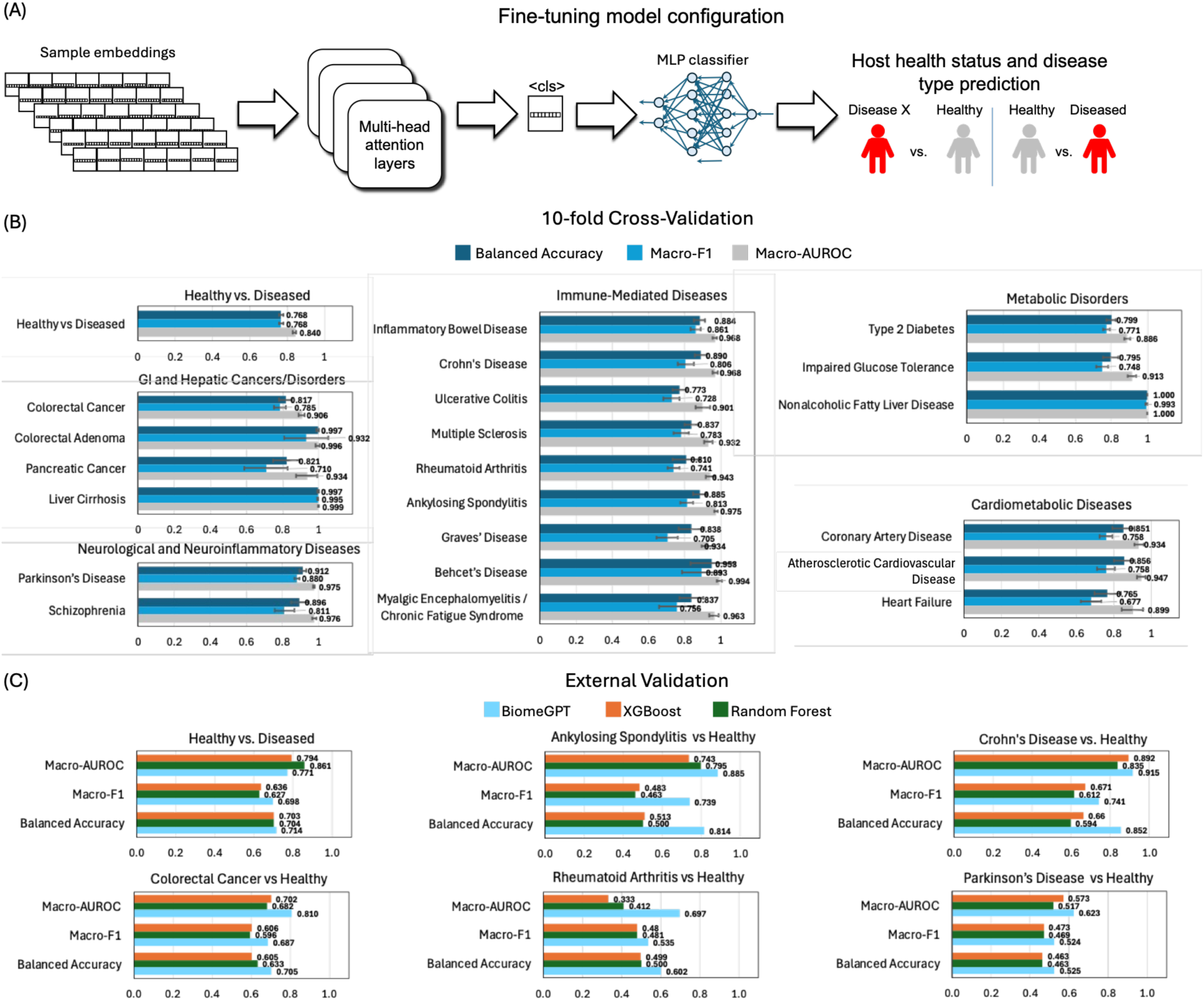
Fine-tuning and evaluation of BiomeGPT for host health status prediction tasks. (A) For all downstream tasks, a classification head was appended to the pretrained model, using the learned sample-level `<cls>` token from the transformer as input. (B) BiomeGPT performance, evaluated by 10-fold cross-validation, for healthy vs diseased and specific-disease vs healthy classification across five major disease categories for diseases with ≥ 20 samples. Crohn’s disease and ulcerative colitis were evaluated separately from the broader IBD. (C) BiomeGPT performance on healthy vs diseased and specific disease vs healthy prediction for five diseases represented in the external validation dataset and comparison with the baseline methods, XGBoost and Random Forest.

#### Distinguishing healthy vs. diseased microbiomes

To assess the capacity of BiomeGPT in distinguishing between healthy and diseased microbiomes, we fine-tuned the pretrained model for Healthy vs. Diseased classification. In 10-fold cross-validation on the training dataset, the model achieved a mean balanced accuracy and macro-F1 score of 0.768 and a macro-AUROC of 0.840 (**Fig. 4B**), suggesting that the model effectively captures broad alterations in microbial composition associated with disease status.

#### Predicting disease type from microbiome profiles across diverse clinical conditions

We next sought to evaluate the model’s capacity to resolve finer-grained diagnostic classes by predicting specific disease types from gut microbiome species-level profiles. This analysis is an initial step toward evaluating whether microbiome representations learned across heterogeneous cohorts could eventually support differential diagnosis by distinguishing among alternative clinical states. To this end, we fine-tuned BiomeGPT across 26 diseases with ≥ 20 samples, training a distinct binary classifier for each disease to differentiate disease-associated from healthy microbiome profiles. The results of this analysis for diseases spanning five major disease categories are shown in **Fig. 4B**. In 10-fold cross-validation, BiomeGPT achieved high performance for all disease types, with mean balanced accuracy of 0.872 ±0.072, mean macro-F1 score of 0.806 ±0.087, and mean macro-AUROC of 0.945 ±0.048 across all diseases. This underscores the model’s capacity to detect intricate disease-specific alterations in microbial composition.

#### External validation across independent host health and disease cohorts

To evaluate the generalizability of BiomeGPT, we performed external validation using an independent dataset comprising 927 samples from studies not included during pretraining and fine-tuning, corresponding to a zero-shot evaluation on previously unseen cohorts. For this evaluation, we retained the same model architecture and initialized all model parameters except those of the classification head from the pretrained BiomeGPT model. All parameters were then updated during fine-tuning using the full training dataset (rather than separately within each cross-validation fold).

For the healthy vs. diseased classification task, the model maintained strong performance, achieving a balanced accuracy of 0.714, a macro F1-score of 0.698, and a macro AUROC of 0.771 (**Fig. 4C**). For the disease-specific prediction tasks (Specific Disease vs. Healthy), BiomeGPT showed the strongest external validation performance for Crohn’s disease, ankylosing spondylitis, and colorectal cancer; however, performance was more modest for rheumatoid arthritis and Parkinson’s disease (**Fig. 4C**). These results show the utility of BiomeGPT in providing reusable pretrained microbiome representations that can be adapted across diverse disease phenotypes and retain predictive utility in independent cohorts, while also highlighting phenotypes for which external cohort alignment may be more challenging.

#### Comparison to baseline methods

To contextualize the external validation performance of BiomeGPT, we benchmarked its fine-tuned classifiers against classical machine learning approaches, including Random Forest and XGBoost, using the same input features and identical external validation cohorts (**Fig. 4C**). For healthy vs. diseased classification, BiomeGPT achieved modestly higher balanced accuracy and a more pronounced improvement in macro-F1 score than XGBoost and Random Forest, although its macro AUROC was lower. In contrast, across all five disease-specific external validation tasks, BiomeGPT outperformed both baseline models in balanced accuracy, macro F1 score, and macro AUROC. The largest improvements were observed for Crohn’s disease and ankylosing spondylitis, while improvements were more modest for Parkinson’s disease.

Overall, BiomeGPT showed its clearest advantage over the conventional machine learning baselines in disease-specific external validation, whereas performance was more comparable for the broader Healthy vs. Diseased task.

### Differential attention analysis to parse phenotype-associated microbial patterns

#### Differential attention patterns across health and disease phenotypes

To investigate which microbial species were differentially emphasized by BiomeGPT between health and disease states, we analyzed attention from the sample-level

`<cls>` token to individual species in the final transformer layer of each fine-tuned classifier. For each species, attention was averaged across heads and separately across healthy and diseased samples, and differential attention was calculated as the mean attention in the diseased group minus that in the healthy group. Positive values therefore indicate greater attention in diseased samples, whereas negative values indicate greater attention in healthy samples. To facilitate interpretation for disease-specific classification tasks, diseases were grouped into five major categories—immune-mediated diseases, GI and hepatic cancers/disorders (non-autoimmune), metabolic disorders, cardiometabolic diseases, and neurological and neuroinflammatory disorders (non-autoimmune)—and species recurrently exhibiting large absolute attention differences across diseases within each category were summarized in **Fig. 5**. Complete differential attention analysis results for individual disease tasks are provided in **Supplementary File 2**.

**Fig. 5.**
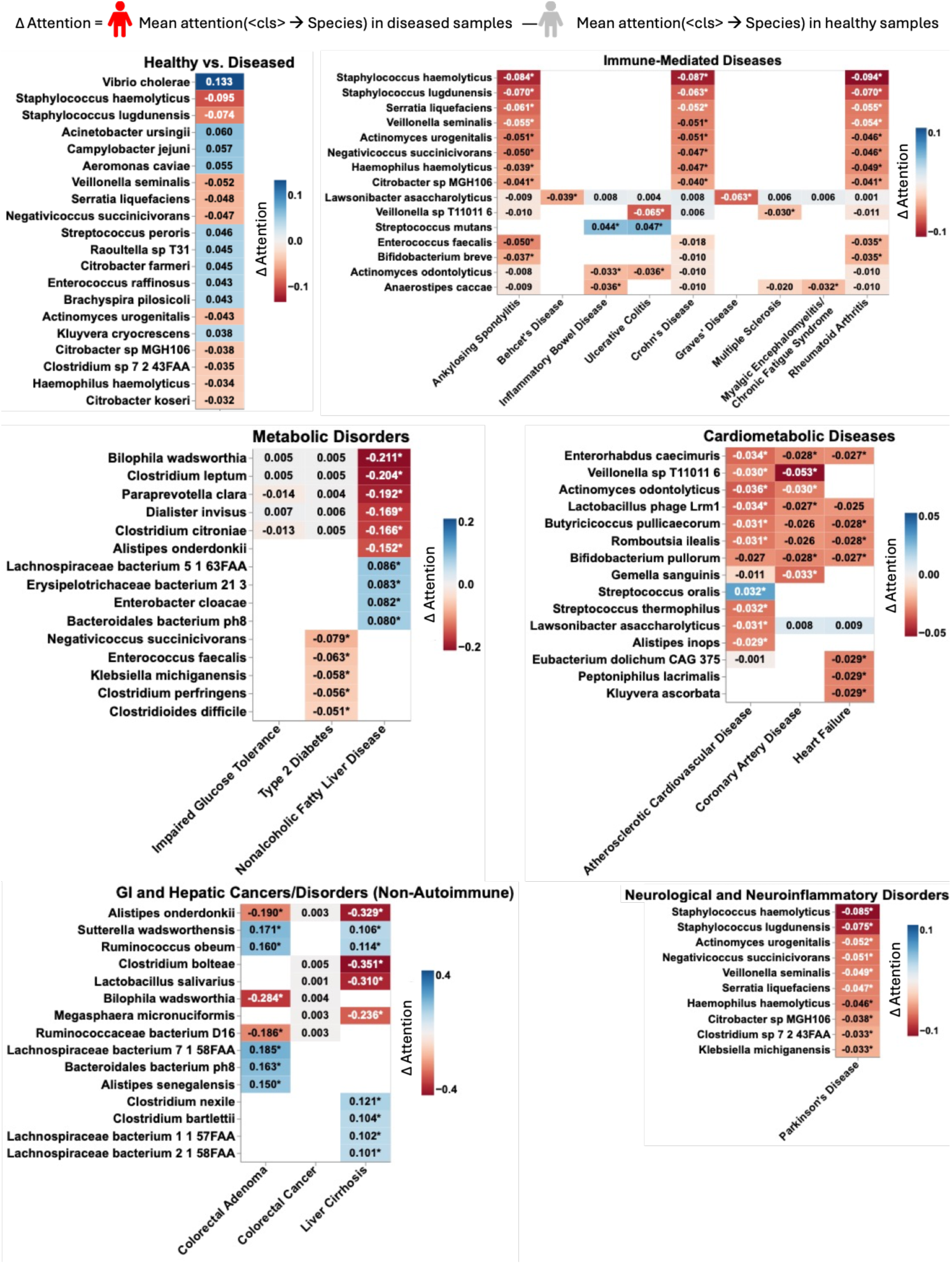
Phenotype-associated microbial signals identified by BiomeGPT differential attention analysis. Differential attention was calculated as the difference in mean attention from the sample-level `<cls>` token to each species between diseased and healthy samples, using attention weights from the final transformer layer averaged across heads. Positive and negative values therefore indicate greater attention in diseased and healthy samples, respectively. For Specific Disease vs. Healthy tasks, diseases were grouped into five major categories. Within each classification task, species were ranked by the absolute magnitude of their differential attention, and the top 10 species were identified. For each disease category, the union of these top 10 species was then ranked by coverage, defined as the number of diseases within the category in which a species appeared among the top 10. Up to 15 species with the highest coverage are shown. Asterisks denote species ranked among the top 10 by absolute differential attention for that disease. White cells indicate |differential attention| < 10⁻³.

For the broad Healthy vs. Diseased classifier, *Vibrio cholerae* showed the largest positive attention difference, followed by *Acinetobacter ursingii*, *Campylobacter jejuni*, and *Aeromonas caviae*. These taxa showed highly restricted distributions in the fine-tuning dataset*: V. cholerae* was detected in 13 of 39 Acute Diarrhea samples and no healthy samples; likewise, *C. jejuni*, *A. caviae*, and *A. ursingii* were detected exclusively in Acute Diarrhea samples. Conversely*, Staphylococcus haemolyticus, Staphylococcus lugdunensis*, and *Veillonella seminalis* showed the strongest negative attention differences and occurred only among healthy samples in the fine-tuning dataset. However, large differential attention shifts were not limited to species with phenotype-restricted distributions. As an example, *Streptococcus lutetiensis* (ranked 33rd; not shown in the figure; see **Supplementary File 2**) was detected in healthy samples and across six disease phenotypes and showed strong positive differential attention despite broadly similar overall prevalence and mean abundance between healthy and diseased samples. These patterns indicate that differential attention reflects phenotype-associated microbial signals that are not determined by species prevalence or abundance alone.

Across disease categories, several species showed recurrent differential attention across multiple related phenotypes. Among immune-mediated diseases, *Lawsonibacter asaccharolyticus* was among the top-10 species by absolute differential attention in two diseases, but also showed non-zero differential attention across all nine diseases in this category; differential attention was positive in six diseases and negative in ankylosing spondylitis, Behçet’s disease, and Graves’ disease, indicating a recurrent but phenotype-dependent attention pattern. Several other species, such as *Staphylococcus haemolyticus*, *Staphylococcus lugdunensis,* likewise showed negative differential attention across several immune-mediated diseases. Recurrent patterns were also evident in metabolic disorders: *Bilophila wadsworthia* and *Clostridium leptum* were among the top 10 species in nonalcoholic fatty liver disease, with strong negative differential attention, while also showing smaller positive shifts in impaired glucose tolerance and type 2 diabetes. In cardiometabolic diseases, the dominant pattern was negative differential attention, with several recurrently selected species receiving greater attention in healthy than in diseased across all diseases (**Fig. 5**).

Beyond these recurrent patterns, several individual diseases showed large disease-specific species attention shifts. The largest such attention shifts were observed among GI and hepatic disorders. For colorectal adenoma, for example, strong positive phenotype-specific shifts included *Lachnospiraceae bacterium* 7 1 58FAA, *Bacteroidales bacterium* ph8, and *Alistipes senegalensis*. In metabolic disorders, nonalcoholic fatty liver disease likewise showed distinct positive differential attention for *Lachnospiraceae bacterium* 5 1 63FAA and *Erysipelotrichaceae bacterium* 21 3.

Together, these analyses revealed both shared and phenotype-specific patterns in the microbial species differentially emphasized by BiomeGPT.

#### Relationship between differential attention and differential abundance

To assess whether phenotype-associated attention shifts are aligned with classical differential abundance signals, we examined the correlation between species-level differential attention and changes in relative abundance across the fine-tuning tasks. For each task (Healthy vs. Diseased and each Specific Disease vs. Healthy comparison), we evaluated the Spearman correlation between differential attention (mean attention in diseased samples minus that in healthy samples) and the log2 fold change in mean abundance between the corresponding groups across species (**Fig. 6**). For most tasks, differential attention was positively correlated with differential abundance. The Healthy vs. Diseased comparison showed a positive correlation (Spearman ρ = 0.66, FDR-adjusted p < 0.05). Particularly strong positive correlations (with Spearman ρ ranging from 0.69 to 0.80) were observed for several disease-specific tasks too, such as Behcet’s disease (ρ = 0.80), IBD (ρ = 0.79), colorectal cancer (ρ = 0.78), rheumatoid arthritis (ρ = 0.76), and type 2 diabetes (ρ = 0.76) (FDR-adjusted p < 0.05 for all; **Fig. 6**). More modest but significant positive correlations were also observed for other diseases such as atherosclerotic cardiovascular disease (ρ = 0.41). However, not all tasks followed this pattern and for several disease-specific comparisons, correlations were weak (0.1 ≤ Spearman ρ < 0.40), near zero (i.e., uncorrelated: |ρ| < 0.1), nonsignificant, or even negative. Specifically, colorectal adenoma (ρ = −0.76) and nonalcoholic fatty liver disease (ρ = −0.59) showed significant negative correlations, whereas liver cirrhosis, Graves’ disease, soil-transmitted helminth infection, acute diarrhea, and *Clostridioides difficile* infection showed no significant correlation after FDR correction. Overall, although differential attention frequently aligns with differential abundance, the variability in the strength and direction of this relationship across tasks suggests that model attention does not simply recapitulate univariate abundance changes.

**Fig. 6.**
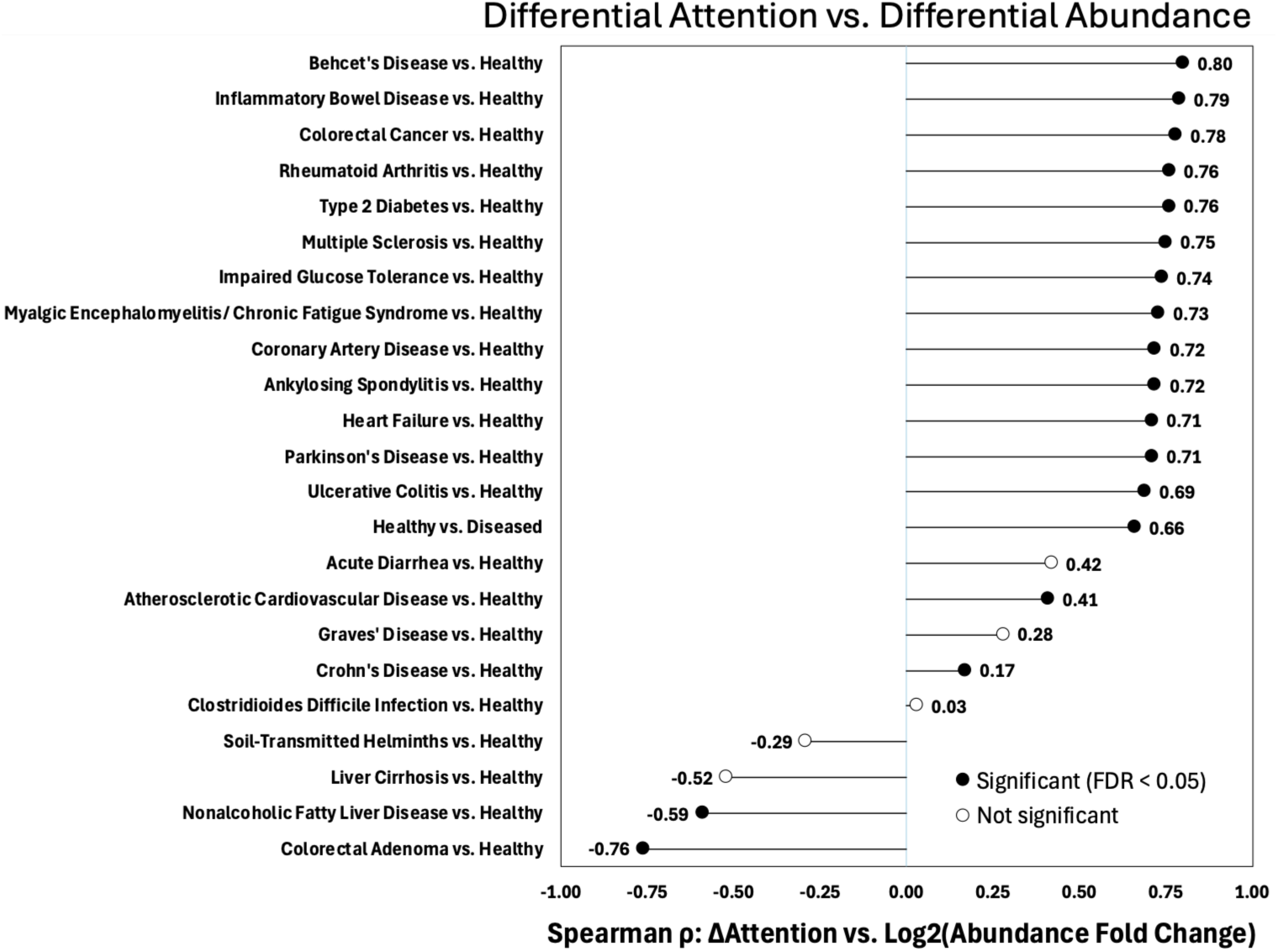
Relationship between differential attention and differential abundance across BiomeGPT fine-tuning tasks. Spearman correlations were evaluated between species-level differential attention and log2 fold change in relative abundance for the Healthy vs. Diseased and Specific Disease vs. Healthy fine-tuning classification tasks. Differential abundance was represented as the log2 fold change in mean relative abundance between diseased and healthy samples.

## Discussion

The growing complexity and scale of microbiome datasets have outpaced the capabilities of traditional analysis pipelines, which remain largely scattered and optimized for a narrow range of task-specific applications. In this study, we introduced BiomeGPT, an abundance- and taxonomy-aware foundation model trained on 13,500 uniformly processed species-level human metagenomic profiles, offering a generalizable framework for microbiome analysis such as microbiome-based host health status prediction. Through large-scale pretraining, BiomeGPT learned representations of microbial community taxonomic composition that could be adapted to distinguish microbiome profiles associated with health and 26 disease states spanning immune-mediated, gastrointestinal, metabolic, cardiometabolic, neurological, and oncological conditions, with predictive utility retained in independent cohorts for several diseases. Differential attention analysis further enabled identification of phenotype-associated microbial signals across diverse disease contexts. These findings establish BiomeGPT as a scalable framework for learning transferable, context-aware representations of human microbiomes and adapting them to diverse downstream prediction tasks.

Within the emerging landscape of microbiome foundation models ^20–22^, BiomeGPT is distinguished by a combination of architectural and representational advances that shape the biological information available to the model during representation learning. By operating directly on species-level profiles rather than profiles at coarser taxonomic resolutions, such as genus level, and explicitly embedding discrete abundance states rather than using abundance-ranked token sequences, BiomeGPT both preserves fine-grained taxonomic resolution and explicitly encodes within-sample abundance patterns in microbial community composition. Importantly, BiomeGPT also incorporates known taxonomic hierarchy directly into the species representation: each species remains a single token, but its embedding integrates learned representations across all eight taxonomic ranks, from Domain through Species. This allows related taxa to share information through their common ancestry while retaining species-specific identity. Consistent with this design, visualization of the learned species representations at higher taxonomic levels (**Fig. 3**) showed that pretraining retained hierarchical taxonomic structure in the embedding space. Notably, the importance of incorporating taxonomic knowledge into microbiome foundation models is also recognized in TaxoMFM ^28^, a contemporaneous model introduced during the preparation of this manuscript. Nevertheless, TaxoMFM incorporates microbial taxonomy through an architecture distinct from that of BiomeGPT; it represents species, genera, families, and orders as separate nodes and encodes their taxonomic relationships through taxonomy-aware attention. In contrast, BiomeGPT encodes the complete eight-rank lineage within each species token while retaining species as the fundamental unit of representation. The two models thus offer distinct but complementary approaches to incorporating taxonomic knowledge into microbiome foundation models.

Building on BiomeGPT’s abundance- and taxonomy-aware input representations, the complementary species- and sample-prompt pretraining objectives further encourage the model to learn cross-species relationships from co-occurrence and context-aware abundance patterns while enabling the sample-level representation to capture broader community structure. Evidence of this broader community structure emerged from the unsupervised evaluation of the learned sample-level representations, which revealed largely distinct regions for gut and non-gut microbiome samples in the embedding space (**Fig. 3**), indicating that the pretrained model captures broad body-site-associated structure without explicit body-site supervision.

An important aspect of microbiome foundation modeling is the ability to transfer information learned across diverse biological and clinical contexts to individual disease phenotypes. Our external validation analyses showed that BiomeGPT had a more consistent advantage over conventional ML models (Random Forest and XGBoost) across all the disease-specific prediction tasks than for the broader Healthy vs. Diseased classification. This may reflect the benefit of broad self-supervised pretraining. For the Healthy vs. Diseased task, conventional ML models are trained on the full heterogeneous disease cohort and therefore have access to much of the same sample diversity as BiomeGPT. In contrast, for individual disease tasks, Random Forest and XGBoost are trained only on samples from the corresponding disease and healthy controls, whereas BiomeGPT draws upon representations learned across a much broader spectrum of microbiome profiles and clinical phenotypes. This broader pretraining may facilitate transfer of community structure and cross-phenotype information across disease contexts that is unavailable to models trained independently for each disease. Nevertheless, reduced cross-cohort transferability across all three models for certain phenotypes, such as Parkinson’s disease (although BiomeGPT still outperformed the baseline models), suggests that cohort alignment and biological heterogeneity remain important determinants of external generalization.

Beyond predictive performance, differential attention provides a model-based view of which microbial species are emphasized differently between health and disease. In the Healthy vs. Diseased classifier, several of the strongest positive differential attention signals corresponded to established or recognized enteric pathogens. For example, *Vibrio cholerae*, which showed the highest positive attention in Healthy vs. Diseased classification is a well-established cause of acute secretory diarrhea and major perturbations of gut community structure ^29, 30^. Other notable species with positive differential attention included *Campylobacter jejuni*, one of the leading causes of bacterial gastroenteritis ^31^; *Aeromonas caviae*, which has been associated with human gastroenteritis and diarrheal disease ^32^; and *Brachyspira pilosicoli*, an enteric spirochete associated with human intestinal spirochetosis and diarrhea ^33^. The convergence of these signals with established disease associations provides biological context for the phenotype-associated patterns captured by differential attention.

Across disease-specific tasks, differential attention revealed both recurrent microbial signals across related phenotypes and shifts specific to individual diseases or disease categories. Several of these patterns also have biological precedent. For example, *Streptococcus mutans* showed positive differential attention in ulcerative colitis and the broader IBD vs. Healthy classification; specific strains of *S. mutans* have previously been associated with ulcerative colitis and shown in mouse models to aggravate colitis ^34^. *Streptococcus oralis* showed positive differential attention in atherosclerotic cardiovascular disease, consistent with a large population study linking gut *S. oralis* and related oral streptococci with subclinical coronary atherosclerosis and systemic inflammatory markers ^35^. *Bilophila wadsworthia* likewise exhibited positive differential attention in type 2 diabetes and impaired glucose tolerance. This is consistent with experimental evidence that *B. wadsworthia* can impair glucose homeostasis in mouse models ^36^ and with a recent study reporting enrichment of the genus *Bilophila* in overweight individuals with early-onset type 2 diabetes ^37^. Interestingly, *B. wadsworthia* showed negative differential attention in nonalcoholic fatty liver disease (NAFLD), consistent with its lower prevalence in NAFLD than in healthy samples in our fine-tuning dataset (34.9% vs. 54.3%), highlighting the phenotype-dependent nature of its differential attention patterns across metabolic disorders.

These concordances support the biological plausibility of several phenotype-associated microbial signals identified by BiomeGPT. Nevertheless, differential attention should be interpreted as a model attribution signal rather than a direct measure of biological importance, and its direction should not be interpreted directly as indicating a beneficial or pathogenic role. Complementary attribution approaches, such as integrated gradients or feature perturbation analyses, could provide an orthogonal assessment of species contributions to model predictions and will be valuable for systematically evaluating the interpretability of BiomeGPT in future studies. A recent work in this direction building on the BiomeGPT architecture explored the use of signed integrated gradients to distinguish species abundance contributions supporting IBD vs. healthy classification ^38^.

The correlation analysis between differential attention and differential abundance (**Fig. 6**) provides additional insight into how BiomeGPT extracts microbiome features across phenotypes. The positive correlations observed for the Healthy vs. Diseased classifier and multiple disease-specific tasks indicate that differential attention frequently tracks phenotype-associated abundance shifts, with species enriched in disease generally receiving greater attention in diseased samples and species enriched in health receiving greater attention in healthy samples. However, this correspondence was far from complete. Even among tasks showing the strongest positive associations, Spearman correlations did not exceed 0.8, while several other disease-specific tasks showed weak, near-zero, nonsignificant, or even negative correlations. Thus, differential attention is not simply equivalent to univariate differential abundance across disease contexts. Rather, the incomplete and heterogeneous correspondence is consistent with BiomeGPT integrating abundance information within a higher-dimensional, context-dependent representation of the microbial community, in which the contribution of a species may depend on its broader community context rather than its marginal abundance change alone. Although the correlation analysis itself does not establish which specific multivariate relationships drive these differences, it suggests that differential attention can provide information complementary to conventional differential abundance analysis rather than merely reproducing it.

While BiomeGPT establishes a strong foundation for general-purpose modeling of the human microbiome, a number of limitations merit consideration. First, performance declined in external evaluations relative to cross-validation and varied across phenotypes, with particularly reduced transferability for some diseases such as Parkinson’s disease. Importantly, this shows that foundation models are not completely immune to the broader challenge faced by AI and ML models when applied to previously unseen cohorts, where distribution shifts can lead to reduced transferability. In microbiome studies, such shifts may arise from differences in cohort composition, geography, clinical characteristics, and other study-specific factors. For Parkinson’s disease, for example, all 490 samples in the training dataset originated from a single study ^39^, which may have limited the diversity of disease-associated variation available during fine-tuning and contributed to reduced transferability to the independent validation cohort. In addition, only five diseases were represented in our external validation dataset, and generalization across the broader set of diseases evaluated in this study remains to be established. Moreover, while uniform taxonomic profiling with MetaPhlAn and quantile-based abundance binning may reduce some technical variation across datasets, they cannot eliminate differences arising from cohort composition, clinical characteristics, or other unmeasured factors. Future studies incorporating geographically and clinically diverse cohorts, broader phenotypic coverage, and more microbiome samples from additional body sites will be important for establishing the generalizability and broader utility of the model. Overall, BiomeGPT represents a step toward a unified, transferable, and reusable representation learning framework for human microbiome research.

## Methods

### Pretraining dataset preprocessing

We excluded samples with incomplete or unclassified phenotypes, as well as those with over 25% unclassified (unknown) taxa from the pretraining dataset. Furthermore, before pretraining, we applied light species-level filtering to remove very rare or low-abundance taxa. Within each phenotype, species were filtered using a minimum prevalence threshold of 1% and a minimum relative abundance threshold of 0.01%. Samples with no species remaining after filtering were excluded, reducing the pretraining dataset from 13,524 to 13,500 samples.

### Taxonomic binning strategy

Prior to model training, we discretized species-level relative abundances into a sample-by-species matrix *A*, where rows represent samples and columns correspond to microbial species. For abundance binning, species with non-zero abundance within each sample were sorted in descending order and divided evenly into *B* bins. The most abundant *N*/*B* species (where *N* is the number of species with non-zero relative abundance in the sample) were assigned to bin *B*, the next *N*/*B* to bin *B* − 1, and so on. Species not detected in each sample (those with zero relative abundance) were assigned a bin value of 0. For samples with fewer than *B* species with non-zero abundance, detected species were distributed proportionally across bins 1 through *B* to ensure that low-diversity samples still spanned the full binned range, avoiding overconcentration of values in the highest bins.

### Taxonomy-aware species representation

To incorporate known microbial taxonomy directly into the model, we represented each species using its complete taxonomic lineage across eight ranks: Domain, Kingdom, Phylum, Class, Order, Family, Genus, and Species. A separate learnable embedding table was defined for each taxonomic rank. For a species *s*, its taxonomic representation was constructed by summing the embedding vectors corresponding to its assigned lineage at all eight ranks:

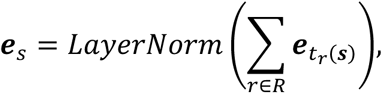

where *R* denotes the set of eight taxonomic ranks (Domain to Species), *t_r_*(*s*) is the taxon assigned to species *s* at rank *r*, ***e****_t_r_(***s***)_* is the learnable embedding vector for that rank, and *e_s_* Is the integrated taxonomy-aware embedding of species *s* Thus, each species remained a single model token, while its representation contained components shared with other species through common taxonomic ancestry as well as a species-specific component. The rank-specific embeddings were learned jointly with the remainder of the model during pretraining.

### Input embeddings and sample representation

Each microbiome sample was represented as a sequence of detected microbial species paired with their corresponding binned abundance values. Species with zero abundance were omitted from the tokenized sequence. For each detected species *s* with abundance bin *b* within a given microbiome sample, the taxonomy-aware species embedding described above was combined with a separate abundance embedding. The scalar abundance bin value was projected through a two-layer multilayer perceptron with a ReLU activation, followed by layer normalization and dropout, to produce an abundance representation *e_b_*. The final input representation for each species-abundance pair was obtained by additive integration of the two embeddings:

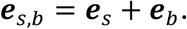

The combined token embeddings were batch normalized before being passed to the transformer encoder.

A special `<cls>` token was also prepended to each microbiome sequence and assigned an abundance bin value of 0. Through transformer attention, the representation at this position was updated using information from the microbial community and it served as the sample-level representation. Consequently, the model represented a microbiome sample through species identities, taxonomic relationships, abundance states, and contextual relationships of its constituent species rather than through their sequence positions.

### Transformer configuration and masked pretraining procedure

BiomeGPT was implemented using transformer encoder layers in `PyTorch`. The model architecture consisted of 8 stacked transformer layers, each containing 8 attention heads and a feedforward sublayer with a hidden dimension of 512. The input to the transformer was the combined taxonomy-aware species and abundance embedding sequence for each sample, with an embedding dimension of 512. Let *E_i_* denote the input embedding sequence for sample *i*. The transformer computed updated representations layer-by-layer as:

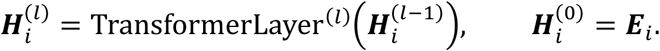

After the final layer, the output at the <cls> position was extracted as the sample-level embedding.

Pretraining used masked abundance prediction with a masking strategy designed for the unordered species composition of microbiome samples. At each epoch, 25% of species with nonzero abundance in each sample were randomly selected for masking, with the masked subset sampled independently across samples and epochs. For masked species, the original abundance bin value was replaced by a dedicated mask value while species identity and its taxonomy-aware representation were retained. Attention was permitted among unmasked species and from masked species to unmasked species and themselves; masked species were prevented from attending to other masked species (**Supplementary Fig. 1**). Samples were padded to a common sequence length within the model input using a dedicated `<pad>` token. Padded positions were excluded from transformer attention using a key padding mask and did not contribute to the reconstruction loss. Species with zero abundance were likewise excluded from the tokenized sequence and loss computation.

BiomeGPT was pretrained using two complementary reconstruction objectives adapted from the gene-prompt and cell-prompt strategies introduced in scGPT ^19^, which we refer to as the *species prompt* and *sample prompt*, respectively. In the species-prompt objective, contextualized species representations produced by the transformer were used to reconstruct the abundance bins of masked species. Each token’s contextual representation was passed through a three-layer MLP decoder with two hidden layers of dimension 512 and LeakyReLU activations, followed by a scalar output predicting the masked abundance bin value. Reconstruction error was computed only at masked positions. The species-prompt objective encouraged the model to infer the binned abundance of a species from the species identities, taxonomic relationships, and abundance context of the other species present in the microbial community. The resulting contextualized `<cls>` representation was then used for the subsequent sample-prompt step. In the sample-prompt objective, the contextualized `<cls>` representation obtained from the species-prompt step was used to replace the initial `<cls>` embedding, and the BiomeGPT transformer was run for a second pass using the same masked species inputs and attention scheme. Masked abundance values were then reconstructed from the resulting contextualized species representations. The species- and sample-prompt reconstruction losses were added together as the loss function.

Pretraining was conducted in two consecutive phases. In the first phase, the model was trained on 13,500 metagenomic samples spanning gut and non-gut body sites. The resulting model was then further pretrained on the gut-specific dataset of 13,333 samples, with model parameters initialized from those learned during the first pretraining phase and subsequently updated during the second phase. For each phase, the corresponding dataset was randomly split into 90% training and 10% internal validation subsets, with the validation subset used to monitor reconstruction performance during training. Pretraining used a batch size of 64 and was performed on a single NVIDIA A100 GPU using Google Colaboratory. Details of model pretraining and pre-trained weights are provided in **Supplementary File 3**.

### Data preprocessing of the fine-tuning dataset

Prior to fine-tuning, all species-level abundance profiles in the gut-specific training dataset underwent preprocessing to enhance gut specificity and reduce noise. First, we excluded species appearing in a predefined blacklist containing 22 microbial species known to be exclusively non-gut or environmental species, representing potential contaminants or sequencing artifacts (see **Supplementary File 1** for the full list). Second, we applied abundance and prevalence filtering on a per-phenotype basis to ensure that retained species were robustly represented within at least one phenotype. Specifically, a species was retained if its relative abundance exceeded 0.1% in at least 2% of samples in one or more phenotypes. The 2% prevalence cutoff was chosen after empirical evaluation of several thresholds, with 2% providing an optimal balance between retaining core gut taxa and excluding low-prevalence or non-gut species. Finally, we filtered borderline species, defined as those belonging to genera that may be occasionally present in the gut transiently or during disease but are also common contaminants originating from skin, oral, or environmental sources (see **Supplementary File 1** for the list of these genera). For each of these genera, all corresponding species detected in our dataset were evaluated across phenotypes (healthy and specific disease). A borderline species was retained within a given phenotype only if it appeared with a relative abundance of ≥0.5% in ≥1% of samples of that phenotype. The higher abundance threshold for borderline taxa reflects their potential contaminant origin, ensuring that only consistently abundant, gut-associated representatives were retained. This multi-tiered filtering strategy aimed to minimize contamination and sparsity while preserving biologically meaningful taxa for fine-tuning classification tasks.

### Fine-tuning model architecture

For all the fine-tuning tasks, the same transformer backbone used in pretraining was retained, and an MLP classification head (two-layer MLP with a 512-dimensional hidden layer, ReLU activation, layer normalization, and a two-unit output layer) was appended to it. The learned sample-level `<cls>` token embedding from the final transformer layer served as input to this classifier. During fine-tuning, the taxonomy-aware species encoder, abundance encoder, and transformer layers were initialized from the pretrained BiomeGPT model, whereas the classification head was initialized separately. All model parameters were subsequently updated during fine-tuning. For each fine-tuning task, we performed feature selection using the least absolute shrinkage and selection operator (LASSO), retaining species with nonzero coefficients as input features. As in pretraining, a subset of species abundance values was masked during training, and the model was jointly optimized for phenotype classification and masked abundance reconstruction.

### Addressing class imbalance for fine-tuning tasks

To address class imbalance in binary disease prediction tasks, we augmented the minority class by generating synthetic samples from existing minority-class profiles. Gaussian noise was added independently to nonzero species abundances, with negative values clipped to zero, and the resulting profiles were renormalized. Synthetic samples were generated until the specified minority-class augmentation target was reached. Augmentation was applied only to the training data; validation and external validation samples were not augmented.

### Differential attention analysis

To quantify phenotype-associated differences in model attention, we extracted attention weights from the final transformer layer of each fine-tuned classifier. For each sample, attention from the `<cls>` query to each species token was averaged across the eight attention heads. Species-level attention was then averaged separately across healthy and diseased samples in which the species was present.

Differential attention for species *s* was calculated as:

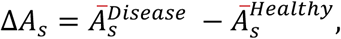

where, *A̅_s_^diseases^* and *A̅_s_^healthy^* denote the mean attention to species *s* averaged across the eight attention heads and across diseased and healthy samples, respectively. Positive differential attention therefore indicates greater attention to a species in diseased samples, whereas negative values indicate greater attention in healthy samples.

### Differential attention-differential abundance correlation analyses

For each fine-tuning task, Spearman’s rank correlation was then calculated across species between differential attention and log₂ fold change in abundance (`scipy.stats.spearmanr`). P values were adjusted across tasks using the Benjamini–Hochberg procedure.

## Code and data availability

All code and data produced in this study, including pretrained weights, are provided within the supplementary materials.

## Author contributions

ARZ conceived the study. NM implemented the model and performed all primary analyses. NY performed the downstream differential attention analysis and analyzed the correlation between differential attention and differential abundance. ARZ and NM analyzed and interpreted the results and drafted the manuscript. All authors have read and approved the final manuscript.

## Acknowledgement

The authors sincerely thank Siyao Zhu for his contributions to the implementation and evaluation of taxonomy-aware microbiome modeling during the earlier stages of this project and Professor Stanislav Dusko Ehrlich for his insightful feedback during the preparation of this manuscript.

