## Supplementary Fig. 1 for "BiomeGPT: A taxonomy-aware foundation model for the human microbiome"

(A)

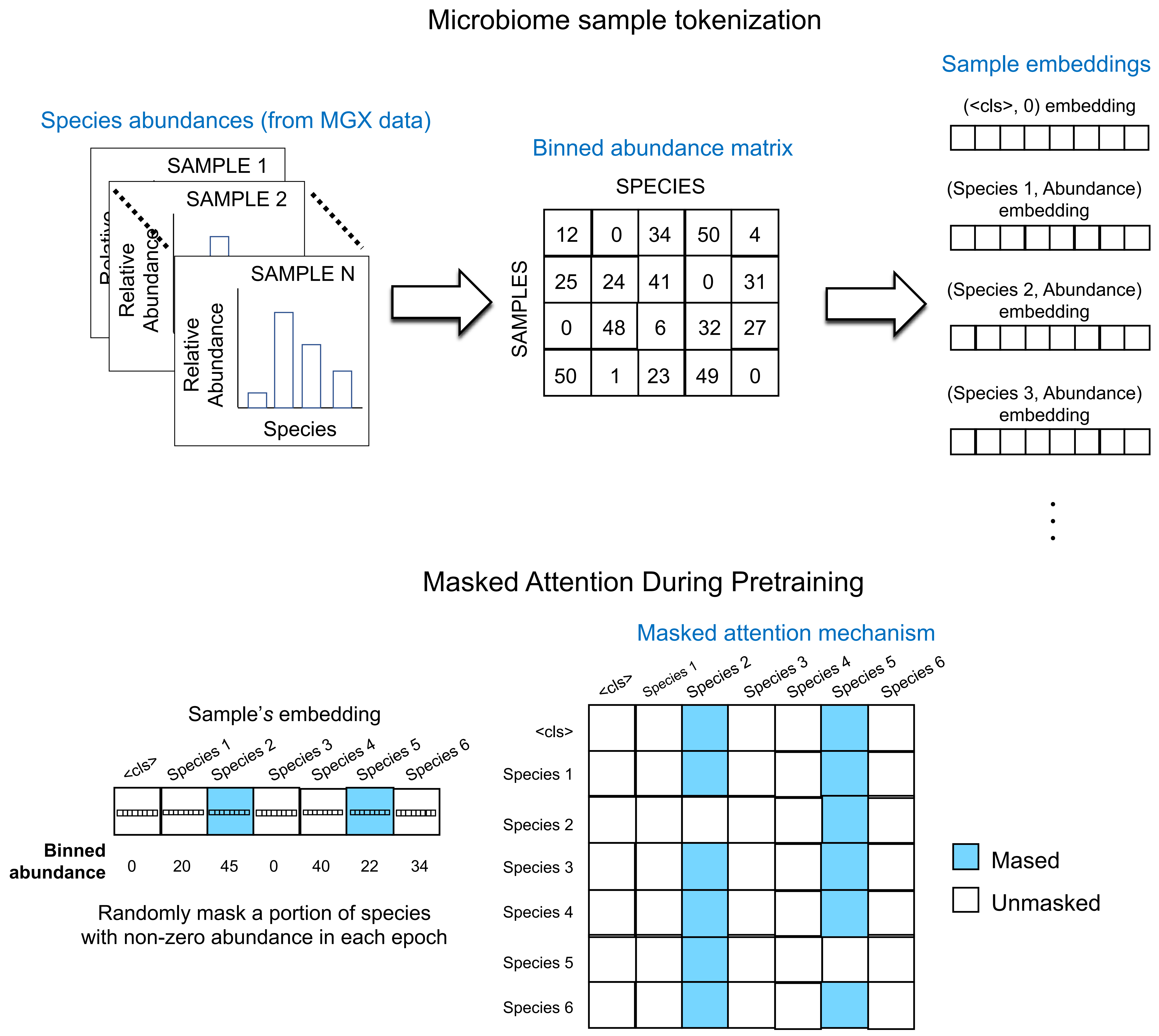


**Supplementary Fig. 1.** Overview of microbiome sample tokenization and masked attention during BiomeGPT pretraining. Species relative abundance profiles derived from metagenomic data are converted into a binned abundance matrix and represented as sample embeddings composed of species and abundance information. During pretraining, a subset of species with nonzero abundance is masked, and the attention mechanism restricts information flow so that masked species attend to the unmasked community context and themselves, while unmasked species attend only to other unmasked tokens.
